# Alcohol Promotes ER+ Breast Cancer Cell Proliferation and Invasion Through the Upregulation of Neuregulin 1-Mediated ER-ErbB3 Crosstalk

**DOI:** 10.64898/2026.09.21.753250

**Authors:** Zhikun Ma, Amanda B Parris, Xiaohe Yang

## Abstract

Alcohol consumption is an established risk factor for breast cancer, with a particularly strong association with estrogen receptor-positive (ER+) disease. Although the estrogenic activity of alcohol is well recognized, how alcohol-induced ER signaling is coupled to growth factor receptor pathways that promote tumor cell growth and progression remains incompletely understood. Here, we identify neuregulin-1 (NRG1) as a functional mediator, linking alcohol-induced ER activity to ErbB3 receptor tyrosine kinase signaling in ER+ breast cancer cells. Under estrogen-depleted conditions, alcohol induced proliferation, clonogenic growth, and S-phase progression in MCF-7 and T47D cells. Alcohol concurrently increased ERα phosphorylation and transcriptional activity, including enhanced ERα occupancy at the endogenous TFF1/pS2 regulatory region, and activated ErbB3 and downstream Akt, ERK, and p38 signaling. Notably, alcohol induced NRG1 expression in both cell lines. Pharmacological inhibition of ER signaling with ICI 182,780 (fulvestrant) substantially attenuated NRG1 induction, ErbB3/RTK activation, and alcohol-promoted growth, indicating that NRG1 induction and engagement of RTK signaling are strongly dependent on functional ER signaling. Conversely, shRNA-mediated NRG1 depletion suppressed alcohol-induced proliferation, clonogenic growth, and cell-cycle progression and markedly reduced alcohol-induced migratory and invasive phenotypes. NRG1 depletion also attenuated ErbB3/Akt/ERK signaling while reducing ERα activation and ERE-dependent transcription, demonstrating a functional contribution of NRG1 to both arms of the signaling response. Together, these findings support a reciprocal ER-NRG1-ErbB3 signaling circuit in which alcohol-induced ER activity promotes NRG1 expression, while NRG1-dependent ErbB3 signaling reinforces ER activity and tumor-promoting phenotypes. Our study identifies NRG1 as a previously unrecognized molecular link between the estrogenic activity of alcohol and growth factor receptor signaling and provides a mechanistic framework for understanding alcohol-associated promotion of ER+ breast cancer.

## Introduction

Breast cancer is the most commonly diagnosed malignancy among women in the United States, and approximately 70% of breast cancers express estrogen receptor α (ERα), a major determinant of tumor development, progression, and therapeutic response^1,2^. Alcohol consumption is a well-established and modifiable risk factor for breast cancer, with epidemiologic studies demonstrating an increased risk even at relatively low levels of consumption^3–6^. Importantly, the association between alcohol consumption and breast cancer is particularly evident for ER-positive (ER+) disease, suggesting that estrogen signaling contributes substantially to alcohol-associated breast cancer risk^7–9^.

Alcohol may influence breast carcinogenesis through multiple mechanisms, including acetaldehyde formation, oxidative stress and DNA damage, and alterations in hormonal signaling^4,10^. The latter is particularly relevant to ER+ breast cancer. Alcohol can increase circulating and tissue estrogen levels and can also enhance ER signaling in breast cancer cells. Under estrogen-depleted conditions, ethanol increases ERα expression and transcriptional activity and promotes proliferation of ER+ breast cancer cells^11–14^. Alcohol has also been reported to activate ERK1/2 and ER-responsive signaling and to attenuate the growth-inhibitory effects of anti-estrogen therapy^15^. Collectively, these findings establish an estrogenic component of alcohol action in ER+ breast cancer. However, how alcohol-induced ER activity is coupled to growth-factor receptor signaling and converted into a broader tumor-promoting response remains incompletely understood.

ER signaling is closely integrated with receptor tyrosine kinase (RTK) pathways in breast cancer. Among the EGFR/ErbB receptor family, ErbB3 (HER3) is particularly relevant because ligand-induced heterodimerization with other ErbB receptors provides a potent signaling platform for PI3K/Akt and MAPK pathways that regulate proliferation, survival, and invasion^16–19^. HER3 signaling has also been implicated in ER+ breast cancer growth and adaptation to endocrine therapy^20,21^. Importantly, ER and ErbB signaling interact bidirectionally. Activation of ErbB-associated kinase pathways can enhance ERα phosphorylation and transcriptional activity, whereas estrogen signaling can alter expression and activity of components of the ErbB network^22–24^. Such reciprocal signaling provides a potential mechanism through which an initial estrogenic stimulus can be amplified through growth-factor pathways.

Neuregulin-1 (NRG1), a major physiological ligand for HER3, is positioned to provide a molecular link between these signaling systems. NRG1 binding promotes formation and activation of HER3-containing receptor complexes and downstream PI3K/Akt and MAPK signaling. NRG1/HER3 signaling has been implicated in breast cancer cell proliferation, survival, motility, metastasis, and resistance to endocrine and HER2-directed therapies^25–28^. Moreover, interactions between NRG1/HER3 and ER signaling have been observed in ER+ breast cancer, particularly in the context of adaptation to estrogen deprivation and endocrine therapy^20,29^. These observations raise an important but unresolved question: does alcohol engage the NRG1/HER3 pathway as a mechanism connecting its estrogenic activity to RTK signaling in ER+ breast cancer cells?

In the present study, we investigated this possibility using MCF-7 and T47D ER+ breast cancer cells under estrogen-depleted conditions. We show that alcohol promotes proliferation and invasive phenotypes while coordinately activating ER and HER3-associated signaling. Alcohol markedly induces NRG1 expression in both cell lines, and this induction is strongly dependent on functional ER signaling. Conversely, depletion of NRG1 attenuates alcohol-induced HER3/RTK signaling, ER activation and transcriptional activity, and the associated proliferative and invasive phenotypes. Together, these findings identify NRG1 as a functional mediator of alcohol-induced ER-HER3 signaling crosstalk and support an ER-NRG1-HER3 signaling circuit through which alcohol can reinforce tumor-promoting signaling in ER+ breast cancer cells.

## Materials and Methods

### Antibodies and reagents

Absolute ethanol (200 proof) was purchased from Thermo Fisher Scientific (Waltham, MA, USA). ICI 182,780 (fulvestrant) was purchased from MedChemExpress (Monmouth Junction, NJ, USA). Primary antibodies against ERα, ERβ, Akt, ERK2, phospho-p38 (Thr180/Tyr182), p38, NRG1, and β-actin were purchased from Santa Cruz Biotechnology (Santa Cruz, CA, USA). Antibodies against phospho-ERα (Ser167), ErbB3, phospho-ErbB3 (Tyr1289), phospho-Akt (Ser473), phospho-ERK1/2 (Thr202/Tyr204), c-Jun, and phospho-c-Jun (Ser73) were purchased from Cell Signaling Technology (Danvers, MA, USA).

### Cell culture and alcohol treatment

Human ER+ breast cancer cell lines MCF-7 and T47D were obtained from the American Type Culture Collection (ATCC; Manassas, VA, USA). Cells were maintained in DMEM/F12 medium supplemented with 10% fetal bovine serum (FBS), 100 units/mL penicillin, and 100 μg/mL streptomycin at 37°C in a humidified atmosphere containing 5% CO₂.

Before alcohol treatment, cells were serum-starved for 24 h unless otherwise indicated. Cells were then treated with the indicated concentrations of ethanol in phenol red-free DMEM/F12 medium supplemented with 5% charcoal-stripped FBS (CS-FBS; Gemini Bio-Products, Sacramento, CA, USA) for the indicated durations. To minimize ethanol evaporation and maintain treatment concentrations, culture dishes were placed inside larger dishes containing ethanol-containing water to establish an ethanol-equilibrated atmosphere.

### Cell proliferation assay

Cell proliferation was assessed using the Cell Proliferation Kit II (XTT; Sigma-Aldrich). Cells were seeded at 1× 10³ cells/well in 96-well plates and cultured for 24 h, followed by serum starvation for 24 h. Cells were then treated with the indicated concentrations of ethanol in phenol red-free DMEM/F12 containing 5% CS-FBS for 5 days. XTT labeling mixture (0.3 mg/mL) was added and cells were incubated for 4 h. Absorbance at 450 nm was measured using a SynergyMx microplate reader (BioTek, Winooski, VT, USA).

### Cell-cycle analysis

Cells were seeded in 60-mm culture dishes and cultured for 24 h. Following serum starvation for 24 h, cells were treated with the indicated concentrations of ethanol in phenol red-free DMEM/F12 containing 5% CS-FBS for 4 h. Cells were collected and fixed overnight in 70% ethanol at −20°C. After washing with PBS, cells were incubated with RNase A (0.5 mg/mL) and propidium iodide (50 μg/mL) for 30 min at 37°C. Cell-cycle distribution was analyzed using a Guava EasyCyte 8 flow cytometer (Millipore, Billerica, MA, USA), and the percentages of cells in G0/G1, S, and G2/M phases were determined using ModFit software.

### Clonogenic assay

Cells were seeded at 1 × 10³ cells/well in 6-well plates and cultured for 24 h. Cells were serum-starved for 48 h and subsequently treated with the indicated concentrations of ethanol in phenol red-free DMEM/F12 containing 5% CS-FBS for 2 weeks. Medium containing the appropriate ethanol concentration was replaced every 3 days. Colonies were stained with 0.5% crystal violet, and colonies containing ≥ 50 cells were counted. Images were acquired using a Nikon SMZ 745T microscope and Nikon Elements Imaging System software.

### Wound-healing assay

Cells were seeded at 2 × 10⁵ cells/well in 6-well plates and cultured in phenol red-free DMEM/F12 containing 5% CS-FBS until approximately 90–100% confluent. A linear wound was generated using a sterile 20 μL pipette tip, and detached cells were removed by washing with PBS. Cells were then cultured in the presence or absence of ethanol for 30 h. Images were acquired at 0 and 30 h. Migration was quantified from the change in wound width and expressed relative to the initial wound width at 0 h. Experiments were performed in triplicate.

### Matrigel invasion assay

Cell invasion was evaluated using Growth Factor Reduced Matrigel Invasion Chambers with 8-μm pore inserts (Corning, NY, USA). Cells (2 × 10⁴) were seeded into the upper chamber in serum-free medium with or without ethanol, while the lower chamber contained 300 μL medium supplemented with 10% FBS as a chemoattractant. After 24 h, cells remaining on the upper surface of the membrane were removed. Cells that had invaded to the lower surface were fixed with methanol for 5 min and stained with crystal violet for 15 min at room temperature. Invaded cells were imaged using a Nikon inverted microscope. Experiments were performed in triplicate.

### Lentiviral NRG1 knockdown

The NRG1 shRNA plasmid (HSH100143) was purchased from GeneCopoeia (Rockville, MD, USA). Control or NRG1 shRNA lentiviral vectors were transfected into 293T cells using the Lenti-Pac HIV Expression Packaging Kit (GeneCopoeia) according to the manufacturer’s instructions. Lentiviral particles were collected at 36 and 48 h after transfection, filtered, and used to infect MCF-7 and T47D cells in the presence of 5 μg/mL polybrene (Sigma-Aldrich, St. Louis, MO, USA) for 24 h. At 48 h after infection, cells were selected with puromycin (1 μg/mL) for 2 weeks to establish stably transduced populations. NRG1 knockdown efficiency was confirmed by Western blotting and qRT-PCR before subsequent experiments.

### ERE luciferase reporter assay

Cells were transfected with an estrogen response element (ERE)-driven luciferase reporter construct using X-tremeGENE 9 DNA transfection reagent (Roche, Indianapolis, IN, USA) according to the manufacturer’s instructions. After 24 h, cells were serum-starved for an additional 24 h and subsequently treated with ethanol and/or ICI 182,780 in phenol red-free DMEM/F12 containing 5% CS-FBS as indicated. Luciferase activity was measured and normalized to Renilla luciferase activity. Results from triplicate transfections were expressed as relative luciferase activity normalized to the corresponding control.

### Western blot analysis

Whole-cell lysates were prepared and protein concentrations were determined using the BCA Protein Assay Kit (Thermo Fisher Scientific). Equal amounts of protein (50 μg) were separated by 10% or 12% SDS-PAGE and transferred to nitrocellulose membranes. Membranes were blocked with 5% nonfat milk for 2 h at room temperature and incubated with the indicated primary antibodies overnight at 4°C. After washing with TBST, membranes were incubated with appropriate horseradish peroxidase-conjugated secondary antibodies for 1.5 h at room temperature. Protein bands were visualized using SuperSignal West Pico chemiluminescent substrate (Thermo Fisher Scientific) and imaged using a FluorChemE imaging system.

### RNA extraction and quantitative real-time PCR

Total RNA was extracted using TRIzol reagent (Life Technologies, Carlsbad, CA, USA) according to the manufacturer’s instructions. Total RNA (1 μg) was reverse transcribed using the iScript cDNA Synthesis Kit (Bio-Rad, CA, USA). Quantitative real-time PCR was performed using All-in-One qPCR Mix (GeneCopoeia) and gene-specific primers in 20-μL reactions. Samples were analyzed in triplicate, and relative gene expression was calculated using the 2^−ΔΔCt^ method with GAPDH as the internal reference.

### Chromatin immunoprecipitation assay

Chromatin immunoprecipitation (ChIP) was performed using the EZ-ChIP Chromatin Immunoprecipitation Kit (Merck Millipore, #17-371) according to the manufacturer’s instructions. Approximately 1 × 10⁷ cells were crosslinked with 1% formaldehyde for 10 min at room temperature, followed by quenching with glycine at a final concentration of 125 mM for 5 min. Cells were lysed, and chromatin was fragmented by sonication to generate DNA fragments of approximately 200–1000 bp.

Diluted chromatin was incubated overnight at 4°C with an ERα antibody or nonspecific IgG control, followed by incubation with protein G agarose for 1 h. Immunocomplexes were sequentially washed, eluted, and reverse-crosslinked at 65°C. Following proteinase K treatment, DNA was purified and analyzed by PCR using primers specific for the TFF1/pS2 regulatory region. PCR products were analyzed by agarose gel electrophoresis to assess ERα occupancy at the TFF1/pS2 locus. ChIP signals were quantified by densitometry and normalized to the corresponding input and/or IgG background. At least three independent biological experiments were performed.

### Statistical analysis

Statistical analyses were performed using GraphPad Prism 10.0 (GraphPad Software, Boston, MA, USA). Data are presented as mean ± SEM. Comparisons between two groups were performed using Student’s *t*-test, and comparisons involving multiple groups were analyzed by analysis of variance (ANOVA), as appropriate. A value of *p* < 0.05 was considered statistically significant; \**p* < 0.05 and \*\**p* < 0.01 are indicated in the figures as appropriate.

## Results

### Alcohol promotes proliferation and cell-cycle progression in ER+ breast cancer cells

We first examined whether alcohol promotes the growth of ER+ breast cancer cells under estrogen-depleted conditions. MCF-7 and T47D cells were cultured in phenol red-free medium containing charcoal-stripped serum and exposed to increasing concentrations of alcohol (0.05–0.2% v/v). XTT analysis showed a dose-dependent increase in viable cell fraction in both cell lines, with the greatest effects observed at 0.2% alcohol. T47D cells exhibited a somewhat stronger response, with significant increases also detected at lower concentrations (Figure 1A).

**Figure 1.**
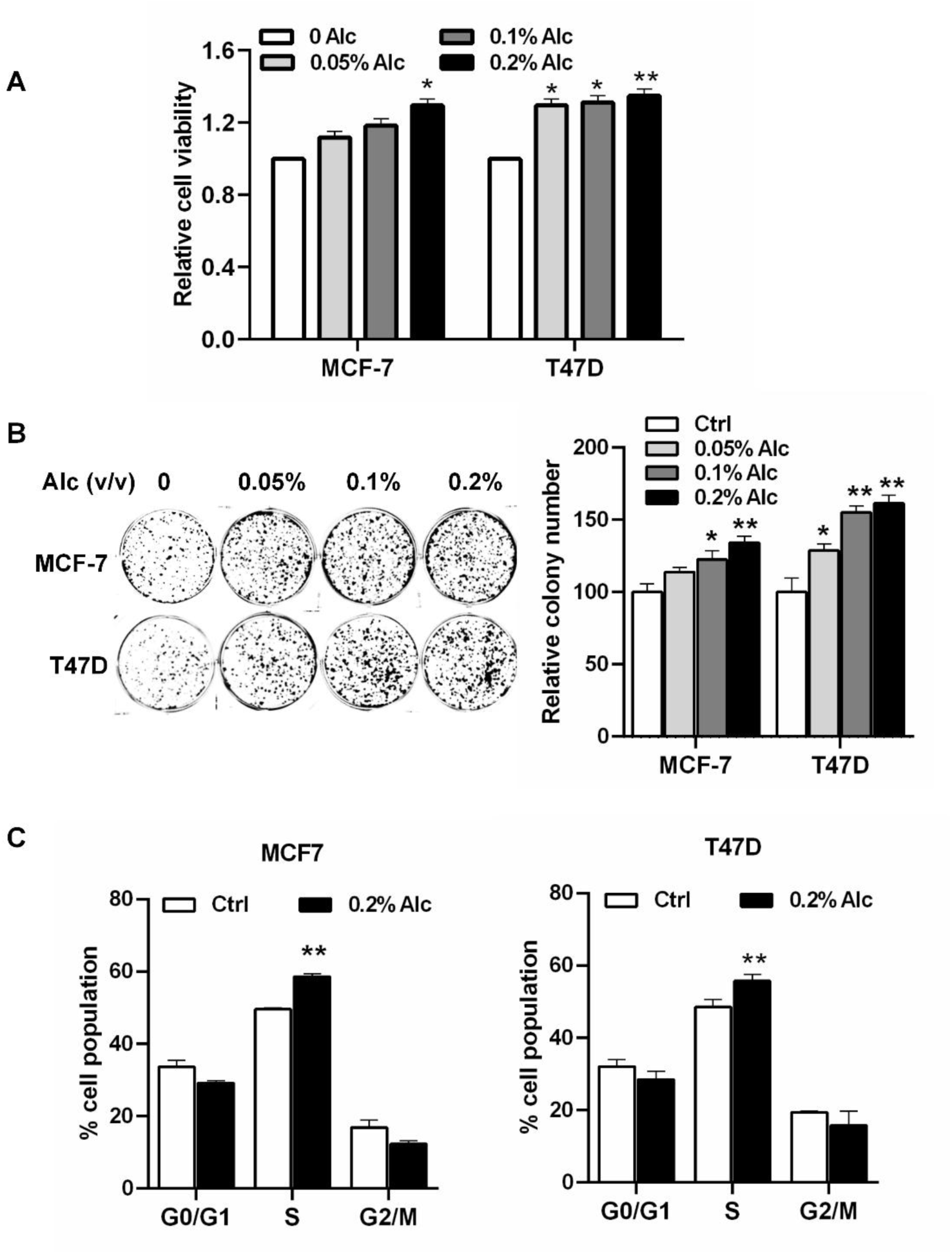
Alcohol promotes proliferation and cell-cycle progression in ER+ breast cancer cells. **(A)** MCF-7 and T47D cells were serum-starved for 24 h and exposed to the indicated concentrations of alcohol (0–0.2% v/v) in phenol red-free DMEM/F12 containing 5% charcoal-stripped FBS for 5 days. Cell viability was assessed by XTT assay and expressed relative to untreated controls. **(B)** MCF-7 and T47D cells were exposed to the indicated concentrations of alcohol for 2 weeks, and clonogenic growth was assessed by crystal violet staining. Representative colony images and quantification relative to untreated controls are shown. **(C)** Serum-starved MCF-7 and T47D cells were treated with 0.2% alcohol for 4 h, and cell-cycle distribution was determined by flow cytometry following propidium iodide staining. Percentages of cells in G0/G1, S, and G2/M phases are shown. Data are presented as mean ± SEM. \**p* < 0.05; \*\**p* < 0.01.

Alcohol similarly enhanced long-term clonogenic growth. Following 2 weeks of exposure, colony formation increased in both MCF-7 and T47D cells in a dose-dependent manner, with a more pronounced response in T47D cells (Figure 1B). These findings indicate that alcohol promotes sustained growth of ER+ breast cancer cells under estrogen-depleted conditions.

We next examined whether this growth response was associated with altered cell-cycle progression. Exposure to 0.2% alcohol significantly increased the proportion of cells in S phase in both cell lines, accompanied by corresponding reductions in the G0/G1 and/or G2/M populations (Figure 1C). Thus, alcohol promotes entry into the DNA-synthetic phase of the cell cycle in ER+ breast cancer cells.

Together, these results demonstrate that alcohol enhances proliferation, clonogenic growth, and S-phase progression under estrogen-depleted conditions. We next examined whether this response was accompanied by activation of ER and receptor tyrosine kinase signaling.

### Alcohol coordinately activates ER and ErbB3/RTK signaling

To define the signaling response to alcohol, MCF-7 and T47D cells were exposed to 0.1–0.2% alcohol for 4 h and analyzed for representative components of ER and RTK pathways. Alcohol increased phosphorylation of ERα and c-Jun in both cell lines, with comparatively modest changes in total ERα, ERβ, and c-Jun protein levels (Figure 2A). In parallel, alcohol increased phosphorylation of ErbB3 and the downstream effectors Akt, ERK1/2, and p38 (Figure 2B), indicating rapid and concurrent activation of ER-associated and ErbB3/RTK signaling.

**Figure 2.**
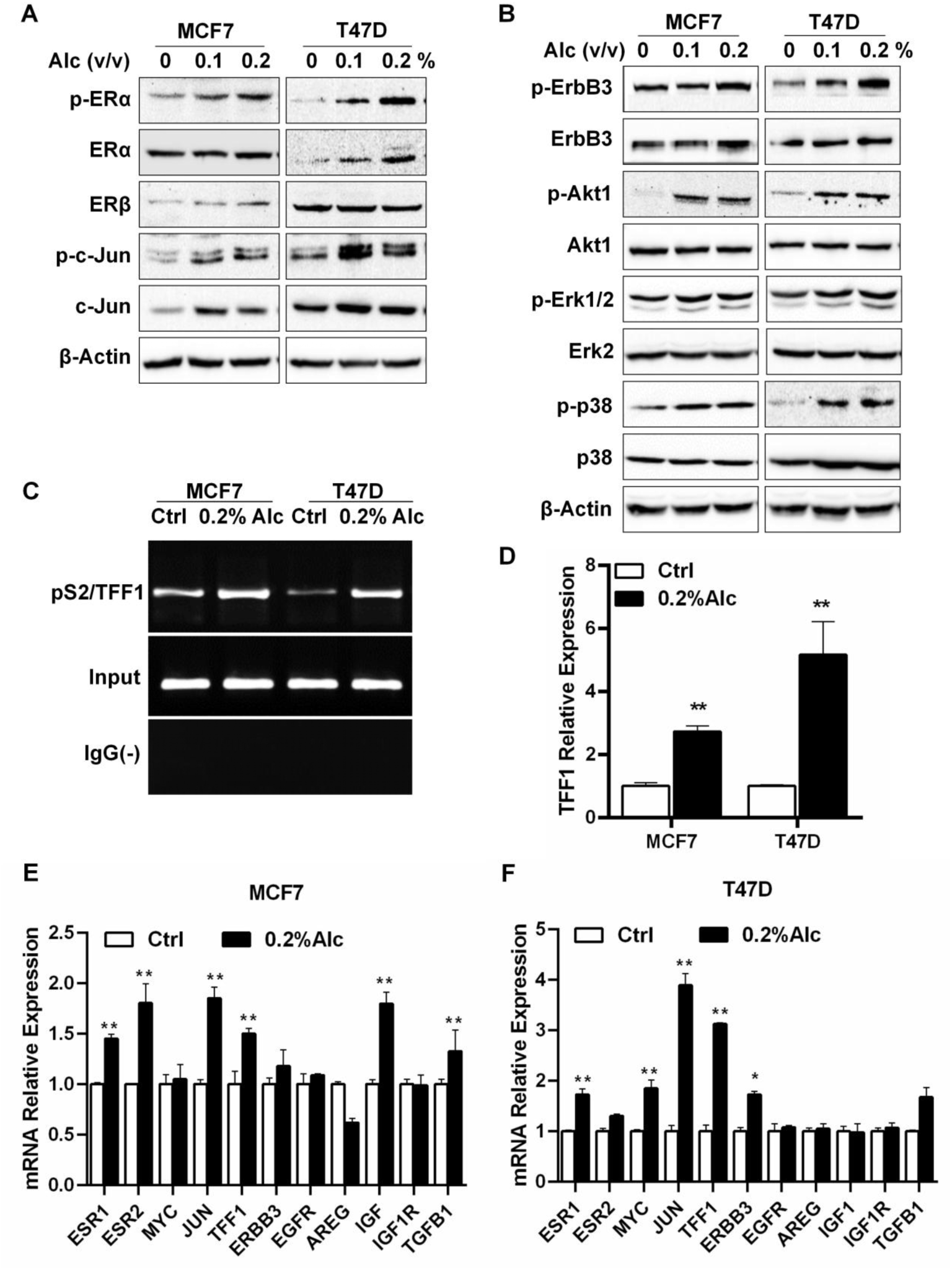
Alcohol coordinately activates ER and ErbB3/RTK signaling in ER+ breast cancer cells. **(A, B)** Serum-starved MCF-7 and T47D cells were exposed to alcohol (0–0.2% v/v) for 4 h, followed by Western blot analysis of the indicated ER-associated **(A)** and ErbB3/RTK-associated **(B)** signaling proteins. β-Actin was used as a loading control. **(C)** Representative ChIP-PCR analysis of ERα occupancy at the TFF1/pS2 regulatory region following treatment with 0.2% alcohol for 4 h. Input chromatin and nonspecific IgG immunoprecipitation are shown as controls. **(D)** Quantification of ERα ChIP-PCR signals normalized to the corresponding input and/or IgG background and expressed relative to untreated controls. **(E, F)** qPCR analysis of the indicated ER- and growth factor signaling-associated genes in MCF-7 and T47D cells following treatment with 0.2% alcohol for 4 h. Expression was normalized to GAPDH and expressed relative to untreated controls. Data are presented as mean ± SEM. \*\**p* < 0.01.

We next assessed whether alcohol-induced ER activation was accompanied by increased ERα engagement at an endogenous target locus. ChIP-PCR analysis showed that 0.2% alcohol markedly increased ERα occupancy at the TFF1/pS2 regulatory region in both MCF-7 and T47D cells (Figure 2C). Quantification demonstrated an approximately 2.5- to 3-fold increase in MCF-7 cells and a greater than 5-fold increase in T47D cells relative to untreated controls (*p* < 0.01; Figure 2D). These data demonstrate that alcohol enhances functional ERα chromatin engagement under estrogen-depleted conditions.

To further characterize the transcriptional response, we examined a panel of genes associated with ER and growth-factor signaling. In MCF-7 cells, alcohol significantly increased expression of several ER-associated genes, including ESR1, ESR2, JUN, and TFF1, whereas most RTK-related genes showed more modest or variable changes (Figure 2E). In T47D cells, significant induction was observed for ESR1, MYC, JUN, and TFF1, while most RTK receptor and ligand transcripts remained comparatively unchanged (Figure 2F). Thus, alcohol elicited a selective transcriptional response rather than a generalized increase in ER- and RTK-associated gene expression.

Collectively, these findings show that alcohol coordinately activates ERα and ErbB3-centered RTK signaling and enhances ER-dependent transcription. This raised the question of whether RTK activation is functionally dependent on ER signaling.

### Alcohol-induced proliferation and ErbB3/RTK signaling are ER-dependent

To determine whether the alcohol response requires ER activity, MCF-7 and T47D cells were treated with the ER antagonist ICI 182,780 (fulvestrant) in the presence or absence of alcohol. Alcohol increased colony formation in both cell lines, whereas ICI reduced basal clonogenic growth and markedly attenuated the alcohol-induced increase (*p* < 0.01; Figure 3A), indicating that ER signaling contributes to the growth-promoting effect of alcohol.

**Figure 3.**
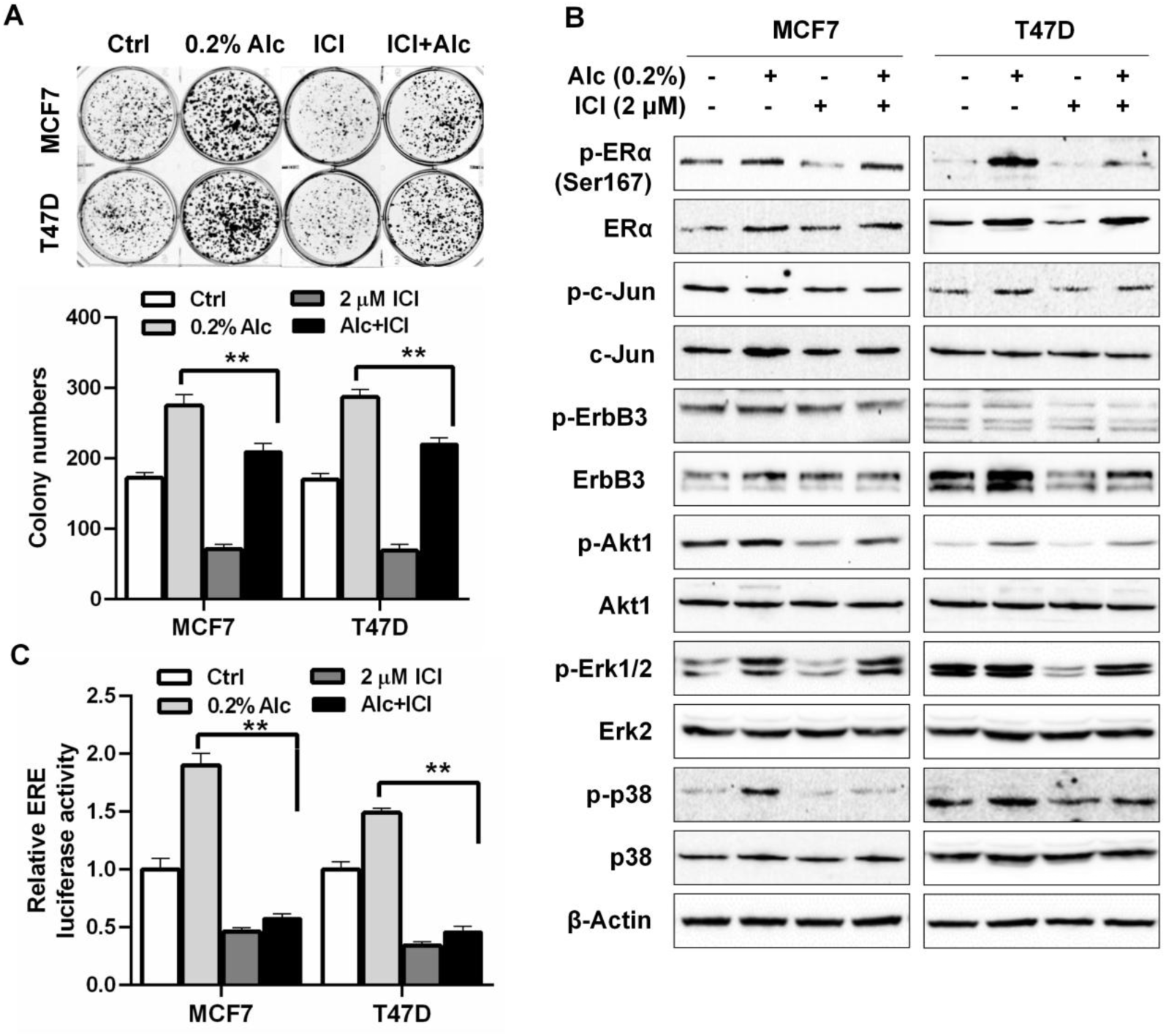
Alcohol-induced proliferation and ErbB3/RTK signaling are ER-dependent. **(A)** MCF-7 and T47D cells were treated with ICI 182,780 (ICI; 2 μM) and/or alcohol (0.2% v/v) for 2 weeks. Colonies were stained with crystal violet, and representative images and colony numbers are shown. **(B)** Serum-starved MCF-7 and T47D cells were treated with ICI (2 μM) for 24 h and/or alcohol (0.2% v/v) for 4 h, followed by Western blot analysis of the indicated ER- and ErbB3/RTK-associated signaling proteins. β-Actin was used as a loading control. **(C)** MCF-7 and T47D cells transfected with an estrogen response element (ERE)-driven luciferase reporter were treated with ICI (2 μM) and/or alcohol (0.2% v/v) as indicated. ER transcriptional activity was determined by luciferase assay and normalized to Renilla luciferase activity. Data are presented as mean ± SEM. ** *p* < 0.01.

We next examined whether ER activity was also required for alcohol-induced RTK signaling. Consistent with Figure 2, alcohol increased phosphorylation of ERα and c-Jun together with ErbB3, Akt, ERK1/2, and p38. ICI reduced ERα expression and attenuated activation of these ER- and RTK-associated signaling components in both cell lines (Figure 3B). These findings indicate that alcohol-induced ErbB3 and downstream RTK signaling depends, at least in part, on functional ER signaling.

Alcohol-induced ER transcriptional activity was further examined using an ERE-driven luciferase reporter. Alcohol significantly increased ERE-luciferase activity in both cell lines, whereas ICI strongly suppressed both basal and alcohol-induced reporter activity (*p* < 0.01; Figure 3C). Together with the increased ERα occupancy at the TFF1/pS2 locus, these findings confirm that alcohol activates ER-mediated transcription under estrogen-depleted conditions.

Taken together, these data establish ER signaling as an upstream component of the alcohol response and suggest that ER activity contributes to the engagement of ErbB3/RTK signaling. Therefore, we next sought to identify a molecular mediator linking these pathways, focusing on NRG1, a major ligand of ErbB3.

### Alcohol induces NRG1 expression in an ER-dependent manner

Having established that ER activity contributes to alcohol-induced ErbB3/RTK signaling, we next examined NRG1 as a potential molecular link between these pathways. Alcohol increased NRG1 protein expression in a dose-dependent manner in both MCF-7 and T47D cells (Figure 4A). Consistent with this response, NRG1 mRNA increased approximately 3-fold in MCF-7 cells and 10-fold in T47D cells following alcohol exposure (*p* < 0.01; Figure 4B). The robust induction of NRG1 in both cell lines, together with concurrent ErbB3 activation, identified NRG1 as a candidate mediator of alcohol-induced ER-RTK signaling.

**Figure 4.**
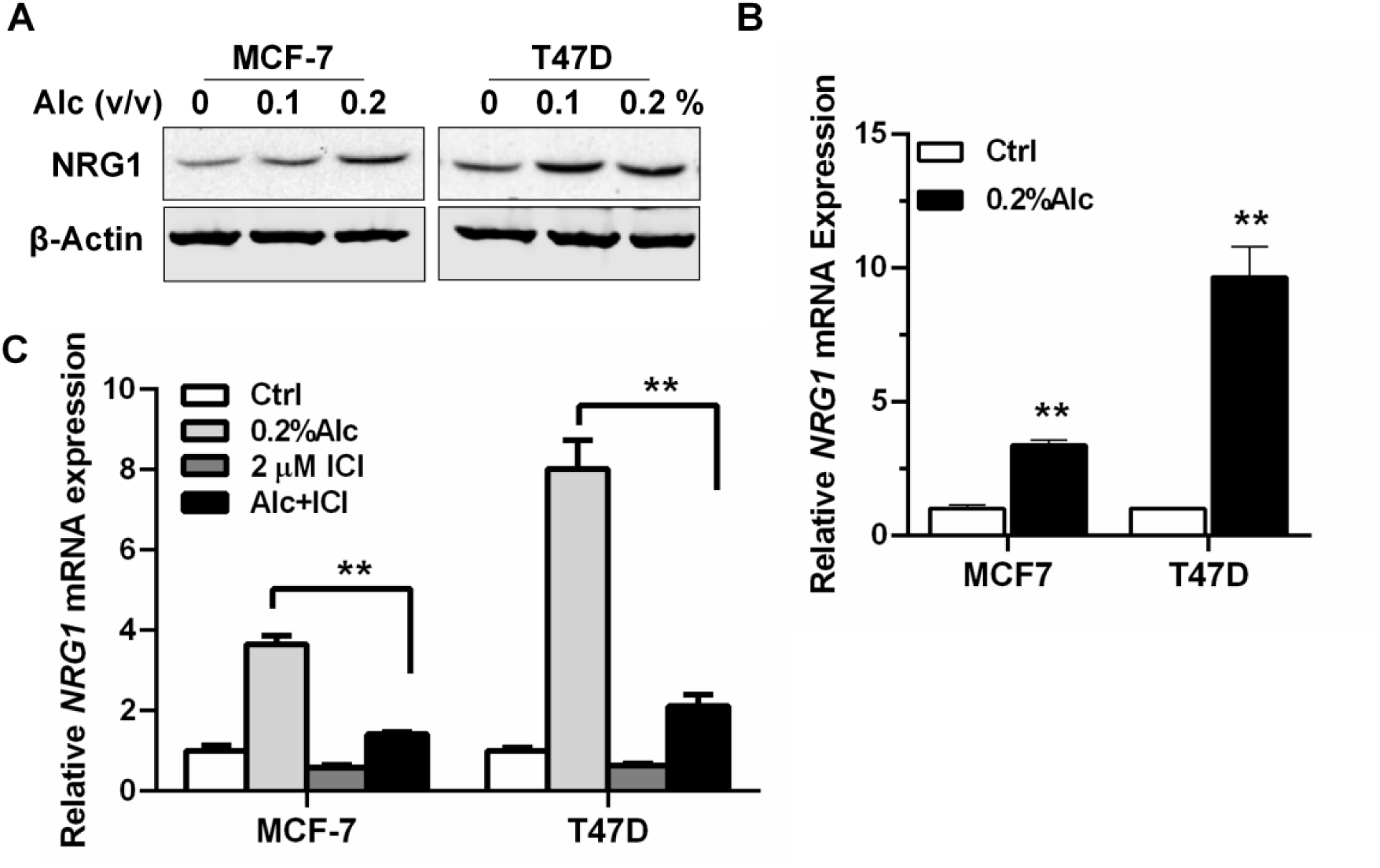
Alcohol induces NRG1 expression in an ER-dependent manner. **(A)** Serum-starved MCF-7 and T47D cells were exposed to the indicated concentrations of alcohol (0–0.2% v/v) for 4 h, followed by Western blot analysis of NRG1. β-Actin was used as a loading control. **(B)** qPCR analysis of NRG1 mRNA expression in MCF-7 and T47D cells following treatment with 0.2% alcohol for 4 h. **(C)** MCF-7 and T47D cells were treated with ICI 182,780 (ICI; 2 μM) and/or alcohol (0.2% v/v), followed by qPCR analysis of NRG1 mRNA expression. Expression was normalized to GAPDH and expressed relative to untreated controls. Data are presented as mean ± SEM. \*\**p* < 0.01.

We next determined whether NRG1 induction depended on ER activity. ICI 182,780 markedly attenuated the alcohol-induced increase in NRG1 mRNA in both MCF-7 and T47D cells (*p* < 0.01; Figure 4C), reducing the response to near-basal levels in MCF-7 cells and substantially suppressing the stronger induction observed in T47D cells. These findings demonstrate that alcohol-induced NRG1 expression is strongly dependent on functional ER signaling.

Thus, NRG1 represents a prominent ER-dependent response to alcohol and a potential link between ER activation and ErbB3 signaling. We therefore next examined whether NRG1 is functionally required for the proliferative response to alcohol.

### NRG1 contributes to alcohol-induced proliferation of ER+ breast cancer cells

To determine the functional role of NRG1, its expression was suppressed by lentiviral shRNA in MCF-7 and T47D cells. Efficient knockdown was confirmed by Western blotting and qPCR, which showed marked reductions in NRG1 protein and mRNA in both cell lines (*p* < 0.01; Figure 5A, B). Alcohol increased the viable cell fraction in control cells, whereas NRG1 depletion markedly reduced cell growth across the alcohol concentrations tested in both MCF-7 and T47D cells (Figure 5C, D). Similarly, alcohol increased clonogenic growth in control cells, while NRG1-depleted cells formed substantially fewer colonies and showed little additional response to alcohol (*p* < 0.01; Figure 5E, F). These findings indicate that NRG1 contributes to both basal growth and the proliferative response to alcohol. NRG1 depletion also altered cell-cycle progression. Alcohol increased the S-phase fraction in control cells, whereas NRG1 knockdown reduced the proportion of cells in S phase in both cell lines (Figure 5G, H). The alcohol-induced shift toward S phase was correspondingly attenuated in NRG1-depleted cells.

**Figure 5.**
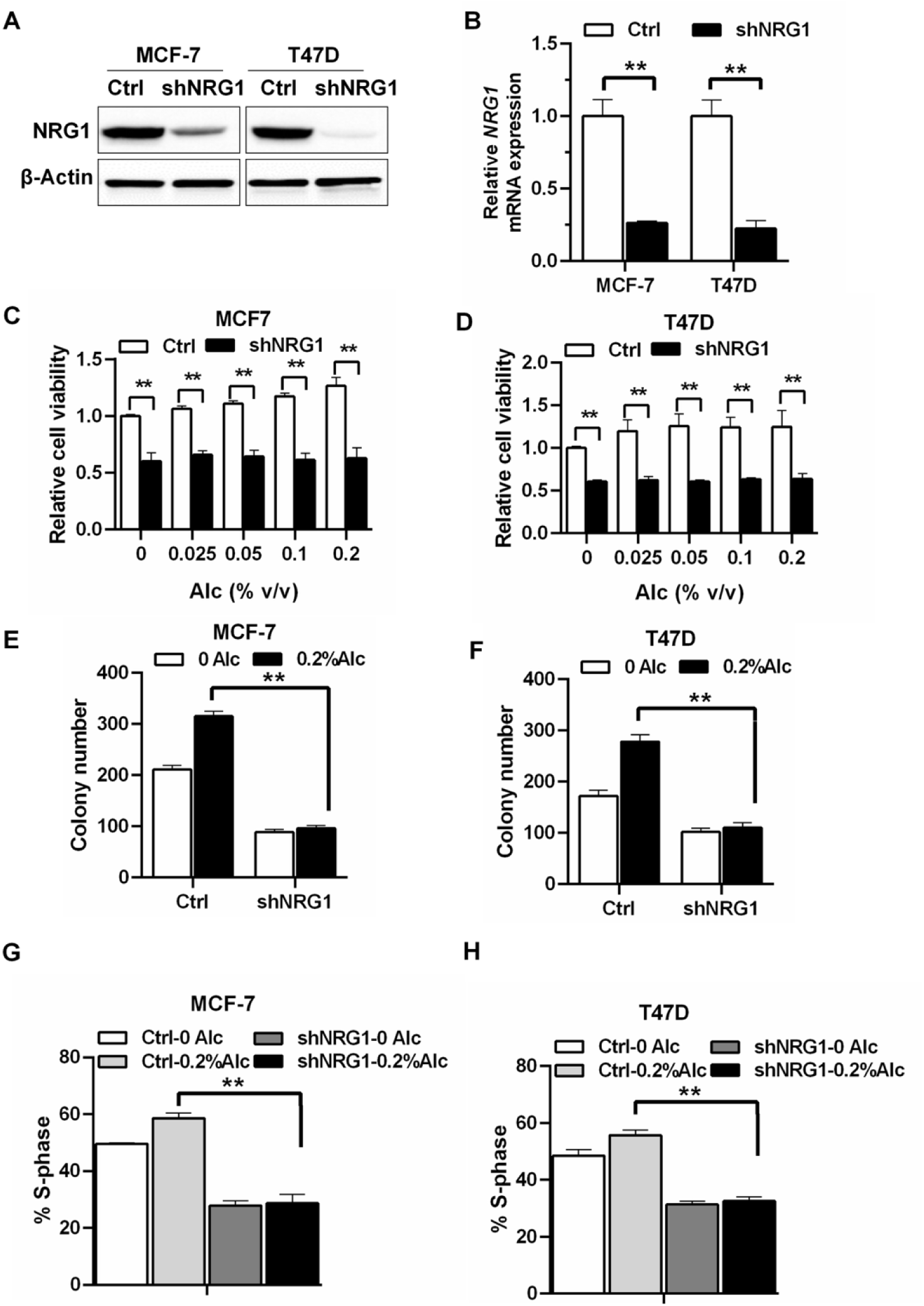
NRG1 contributes to alcohol-induced proliferation and cell-cycle progression in ER+ breast cancer cells. **(A, B)** NRG1 knockdown in stably transduced shControl and shNRG1 MCF-7 and T47D cells was confirmed by Western blot analysis **(A)** and qPCR **(B)**. β-Actin was used as a loading control for Western blotting, and NRG1 mRNA expression was normalized to GAPDH. **(C, D)** shControl and shNRG1 MCF-7 and T47D cells were exposed to the indicated concentrations of alcohol (0–0.2% v/v) for 5 days, followed by XTT assay. Cell viability was expressed relative to the corresponding untreated shControl cells. **(E, F)** shControl and shNRG1 MCF-7 and T47D cells were treated with or without 0.2% alcohol for 2 weeks, and clonogenic growth was quantified following crystal violet staining. **(G, H)** Serum-starved shControl and shNRG1 MCF-7 (G) and T47D (H) cells were treated with or without 0.2% alcohol for 4 h. Cell-cycle distribution was determined by flow cytometry, and the percentage of cells in S phase is shown. Data are presented as mean ± SEM. \*\**p* < 0.01.

Together, these results establish a functional role for NRG1 in alcohol-promoted proliferation, clonogenic growth, and cell-cycle progression. We next examined whether NRG1 also contributes to alcohol-induced migratory and invasive phenotypes.

### NRG1 mediates alcohol-induced migration and invasion of ER+ breast cancer cells

In Matrigel invasion assays, alcohol significantly increased invasion of control MCF-7 and T47D cells (Figure 6A, B). NRG1 knockdown markedly reduced basal invasion and substantially attenuated the increase associated with alcohol treatment in both cell lines (*p* < 0.01), indicating that NRG1 contributes to alcohol-promoted invasive activity. The effect of NRG1 depletion on cell migration was further examined using a wound-healing assay in MCF-7 cells. Alcohol enhanced wound closure in control cells, whereas NRG1 knockdown reduced basal migration and markedly diminished the migratory response to alcohol (*p* < 0.01; Figure 6C). These findings were consistent with the invasion assays and further support a role for NRG1 in alcohol-induced cell motility. Thus, the functional contribution of NRG1 extends beyond proliferation to migratory and invasive phenotypes. We then proceeded to examine whether NRG1 is required for the coordinated activation of ER and ErbB3/RTK signaling.

**Figure 6.**
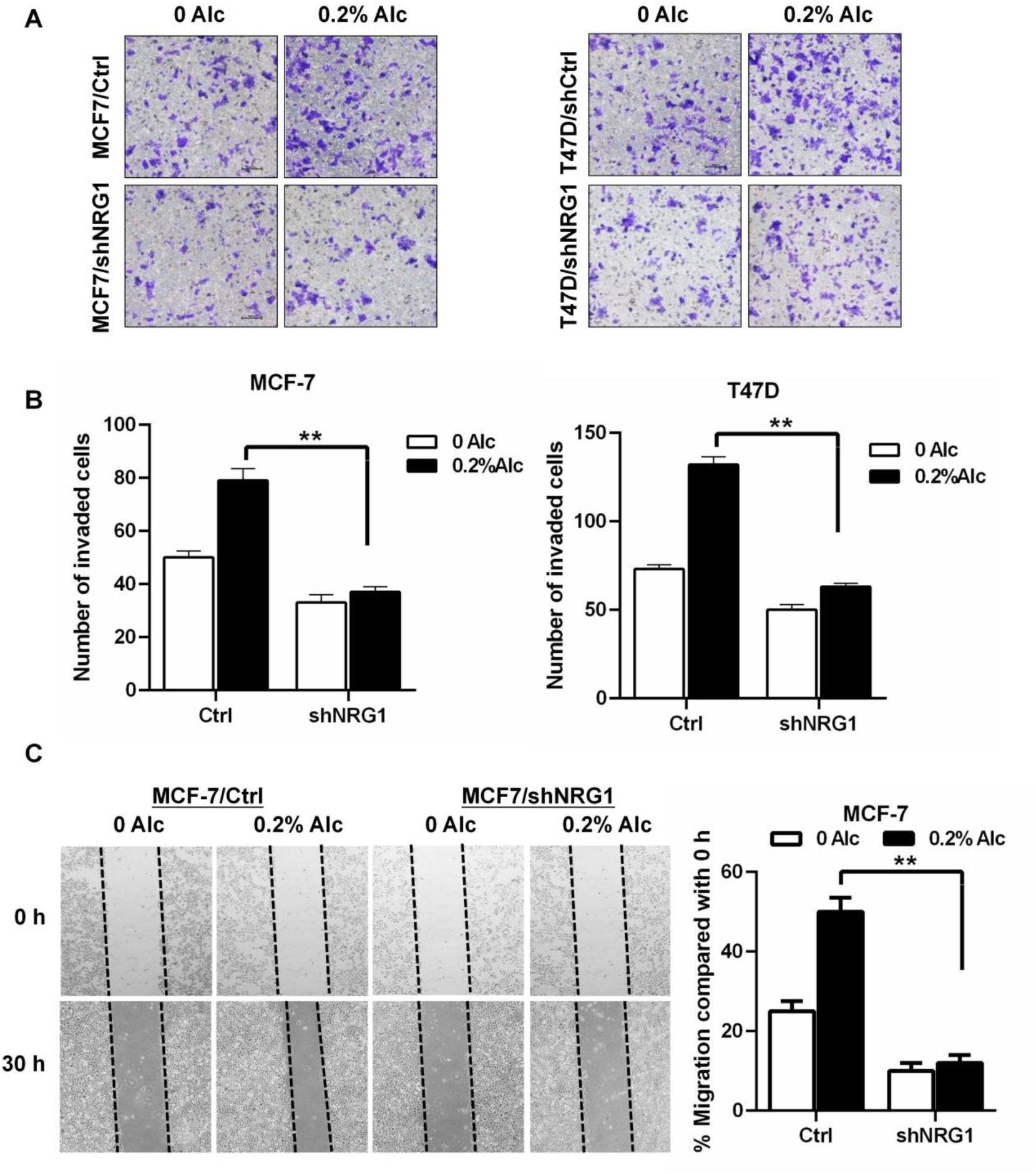
NRG1 mediates alcohol-induced migration and invasion of ER+ breast cancer cells. **(A, B)** shControl and shNRG1 MCF-7 and T47D cells were treated with or without 0.2% alcohol for 24 h and analyzed using Matrigel invasion assays. Representative images of crystal violet-stained cells that invaded through the membrane are shown in **(A)**, and numbers of invaded cells are quantified in **(B). (C)** Cell migration was assessed by wound-healing assay in shControl and shNRG1 MCF-7 cells treated with or without 0.2% alcohol. Representative images were acquired at 0 and 30 h; dashed lines indicate wound boundaries. Migration was quantified from changes in wound width relative to 0 h. Data are presented as mean ± SEM. \*\**p* < 0.01.

### NRG1 mediates alcohol-induced ER–ErbB3 signaling crosstalk

To determine whether NRG1 participates in the signaling response to alcohol, control and NRG1-depleted MCF-7 and T47D cells were analyzed following alcohol exposure. In control cells, alcohol increased NRG1 expression together with phosphorylation of ERα and c-Jun. These responses were markedly attenuated following NRG1 knockdown (Figure 7A), indicating that NRG1 contributes to alcohol-induced ER pathway activation. NRG1 depletion similarly impaired ErbB3-associated signaling. Alcohol increased phosphorylation of ErbB3 and the downstream effectors Akt and ERK1/2 in control cells, whereas these responses were substantially diminished in NRG1-depleted cells (Figure 7B). These findings establish NRG1 as an important mediator of alcohol-induced ErbB3/RTK signaling.

**Figure 7.**
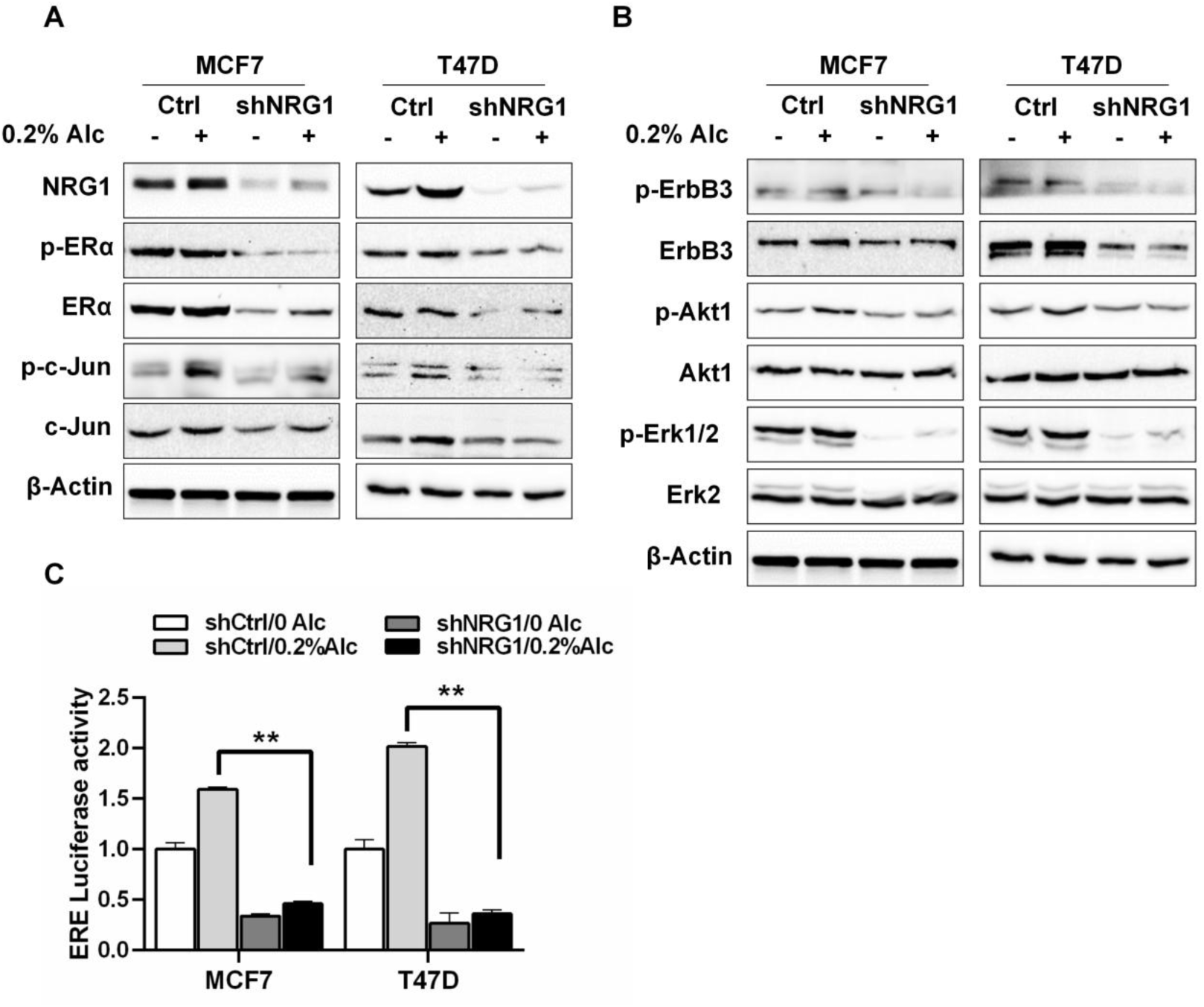
NRG1 mediates alcohol-induced ER–ErbB3 signaling crosstalk. **(A, B)** Serum-starved shControl and shNRG1 MCF-7 and T47D cells were treated with or without 0.2% alcohol for 4 h, followed by Western blot analysis of NRG1 and the indicated ER-associated **(A)** and ErbB3/RTK-associated **(B)** signaling proteins. β-Actin was used as a loading control. **(C)** shControl and shNRG1 MCF-7 and T47D cells were transfected with an ERE-driven luciferase reporter, serum-starved for 24 h, and subsequently treated with or without 0.2% alcohol for 4 h. ER transcriptional activity was determined by luciferase assay and normalized to Renilla luciferase activity. Data are presented as mean ± SEM. \*\**p* < 0.01.

We next examined whether NRG1 also contributes to ER transcriptional activity. Alcohol significantly increased ERE-driven luciferase activity in control MCF-7 and T47D cells, whereas NRG1 knockdown strongly reduced both basal and alcohol-induced reporter activity (*p* < 0.01; Figure 7C). Thus, NRG1 is required for full ER transcriptional activity in response to alcohol. Together with the ER-dependent induction of NRG1 demonstrated in Figure 4, these findings support a reciprocal relationship in which alcohol-induced ER activity promotes NRG1 expression, while NRG1-dependent ErbB3 signaling reinforces ER activation and transcriptional activity. This ER-NRG1-ErbB3 crosstalk provides a mechanistic link between alcohol exposure and the proliferative and invasive phenotypes observed in ER+ breast cancer cells.

## Discussion

Alcohol consumption is an established risk factor for breast cancer, with epidemiologic studies demonstrating a stronger association with ER-positive than with ER-negative disease^8^. Experimental studies have provided biological support for this association by showing that alcohol can enhance estrogen availability and activate ER signaling. However, ER functions within an interconnected signaling network, and the mechanisms through which the estrogenic effects of alcohol are coupled with growth factor receptor pathways remain incompletely understood. In the present study, we identify NRG1 as an important molecular mediator linking alcohol-induced ER activity with ErbB3/RTK signaling in ER+ breast cancer cells. Alcohol promoted ER transcriptional activity and markedly induced NRG1 expression, whereas inhibition of ER signaling substantially attenuated NRG1 induction. Conversely, depletion of NRG1 impaired alcohol-induced ErbB3 and downstream signaling as well as ER activation and transcriptional activity, accompanied by suppression of proliferative and invasive phenotypes. Collectively, these findings support an ER-NRG1-ErbB3 signaling circuit through which alcohol may amplify growth-promoting signaling in ER+ breast cancer cells.

The present findings extend previous studies of the estrogenic activity of alcohol. Alcohol-associated activation of ER signaling has been attributed to several mechanisms, including changes in estrogen metabolism and availability, modulation of ERα expression, and activation of signaling pathways capable of regulating ER activity. Our findings confirm that alcohol can activate ER signaling even under estrogen-depleted culture conditions. Importantly, increased ERα phosphorylation was accompanied by increased ERE reporter activity and enhanced ERα occupancy at the endogenous estrogen-responsive TFF1/pS2 locus. Thus, the response is not limited to biochemical phosphorylation of the receptor but extends to functional engagement of ER-dependent transcription. These observations provide a mechanistic context for the preferential association between alcohol exposure and ER+ breast cancer and reinforce the concept that ER signaling represents an important component of the biological response of breast cancer cells to alcohol.

At the same time, the present results suggest that ER activation alone does not fully account for the growth-promoting effects of alcohol. A major finding of this study was the prominent induction of NRG1, a principal ligand of ErbB3, in both MCF-7 and T47D cells. This observation is particularly relevant because ErbB3 occupies an unusual position within the ErbB receptor network. Although its intrinsic kinase activity is limited, ligand-activated ErbB3 forms signaling-competent heterodimers with other ErbB receptors and provides a particularly effective platform for activation of PI3K/Akt, while ErbB receptor complexes also engage MAPK/ERK signaling^30^. These pathways regulate proliferation and survival and can also modify ER function. NRG1 induction therefore provides a biologically plausible mechanism by which an alcohol-induced estrogenic signal could be propagated into a broader growth-factor signaling response.

The relationship between ER and ErbB signaling is well recognized in breast cancer. Rather than operating as independent pathways, ER and growth-factor receptor signaling can interact bidirectionally^24^. ErbB-associated PI3K/Akt and MAPK signaling can regulate ER phosphorylation and transcriptional activity, while ER-dependent transcription can alter expression of growth factors and components of RTK signaling networks. Such reciprocal interactions have been particularly implicated in tumor progression and adaptation to endocrine therapy^24^. Our findings place NRG1 at an important point within this interaction in the context of alcohol exposure. Fulvestrant strongly attenuated alcohol-induced NRG1 expression together with ErbB3/RTK activation, indicating that functional ER signaling lies upstream of much of the NRG1 response. Conversely, NRG1 depletion suppressed ErbB3/Akt/ERK signaling and reduced ER phosphorylation and ERE-dependent transcription. Taken together, these reciprocal perturbation experiments argue that NRG1 is not simply another alcohol-responsive gene but participates functionally in communication between the ER and ErbB3 pathways.

This observation leads to an important conceptual implication of the study. Alcohol may act less by creating a novel signaling pathway than by engaging and reinforcing an existing ER-growth factor signaling network. We propose a model in which alcohol initially enhances ER activity, which promotes NRG1 expression through an ER-dependent mechanism. Increased NRG1 then facilitates ErbB3 signaling and activation of downstream pathways including Akt and ERK, which are themselves capable of reinforcing ER activity. Such reciprocal signaling provides a potential amplification mechanism whereby an initial estrogenic response to alcohol can be translated into a more sustained mitogenic program. The present experiments do not establish that ERα directly binds regulatory elements of the NRG1 gene, nor do they demonstrate that extracellular NRG1 establishes a classical autocrine loop. Therefore, the precise transcriptional and ligand-receptor steps remain to be defined. Nevertheless, the complementary effects of ER inhibition and NRG1 depletion provide functional evidence for a reciprocal ER-NRG1-ErbB3 relationship.

The biological consequences of NRG1 depletion further support this model. Suppression of NRG1 reduced basal growth as well as the additional proliferative response to alcohol, suggesting that alcohol exploits a signaling pathway already important for the growth of ER+ breast cancer cells rather than generating an entirely alcohol-specific pathway. NRG1 depletion also altered cell-cycle progression and markedly reduced clonogenic growth. Importantly, the consequences extended beyond proliferation: NRG1 depletion strongly attenuated the invasive response to alcohol and was accompanied by reduced migratory activity. NRG1/ErbB3 signaling has been implicated more broadly in breast cancer cell survival, motility, progression, and therapeutic resistance^31–33^. Recruitment of this pathway by alcohol therefore provides a potential mechanistic connection between alcohol exposure and multiple tumor-promoting cellular behaviors rather than proliferation alone.

These findings also provide an interesting perspective on endocrine therapy. Cross-talk between ER and growth-factor receptor pathways is an established mechanism through which ER+ breast cancer cells adapt to estrogen deprivation or ER-directed therapy, and NRG1/ErbB3 signaling has been implicated in this adaptive process. The ability of alcohol to stimulate ER activity while simultaneously engaging NRG1-ErbB3 signaling therefore raises the possibility that alcohol exposure could influence cellular responses to endocrine treatment. This possibility is particularly intriguing because the same reciprocal signaling that promotes growth under estrogen-depleted conditions could potentially provide alternative proliferative signals when ER signaling is therapeutically constrained. Our current experiments were not designed to test endocrine resistance, and this implication should therefore remain a hypothesis. Nevertheless, determining whether alcohol-induced NRG1/ErbB3 signaling modifies sensitivity to anti-estrogen or aromatase-directed therapies represents an important direction for future investigation.

The identification of NRG1 as a functional mediator also has potential therapeutic implications. Unlike the broad biological effects of alcohol, the NRG1-ErbB3 interaction represents a defined signaling node that is potentially amenable to therapeutic intervention. ErbB3 has increasingly been investigated as a therapeutic target because of its role in PI3K/Akt signaling and resistance to ER- and ErbB-directed therapies. Our observation that NRG1 depletion simultaneously attenuated alcohol-induced proliferation, invasion, RTK signaling, and ER transcriptional activity suggests that disruption of this axis can interfere with several components of the alcohol response. These results do not establish NRG1 or ErbB3 inhibition as a strategy for preventing alcohol-associated breast cancer, but they identify the pathway as a mechanistically relevant candidate for further investigation, particularly in the setting of ER+ disease and endocrine therapy.

The findings should also be considered within the broader biology of alcohol-associated breast cancer. Alcohol exposure can affect mammary carcinogenesis through multiple mechanisms, including systemic hormonal changes, acetaldehyde-mediated DNA damage, oxidative stress, altered one-carbon metabolism, and effects on the tissue microenvironment. These mechanisms are not mutually exclusive and may differ according to exposure level and duration. In particular, higher or prolonged alcohol exposure can produce cellular stress and genotoxic effects that differ from the proliferative signaling examined here. The ER-NRG1-ErbB3 mechanism identified in this study should therefore be viewed as one component of the biological response to alcohol; one that may be especially relevant to ER-responsive cells and to exposure conditions favoring proliferative signaling rather than overt toxicity. This distinction may help reconcile the estrogenic and mitogenic effects observed at lower experimental exposures with the DNA-damaging and growth-inhibitory consequences reported under more intensive exposure conditions.

Our mechanistic work used two established ER+ breast cancer cell lines, so it cannot capture systemic alcohol metabolism, endocrine signaling, stromal crosstalk, or in vivo exposure dynamics. The ethanol concentrations we used were chosen to define pathway responses under controlled conditions and should not be equated with blood alcohol levels or human drinking behavior. While pharmacological ER inhibition shows that NRG1 induction is strongly ER-dependent, how ER regulates NRG1 remains unclear; determining whether this is direct or indirect will require profiling ER occupancy and regulatory elements at the NRG1 locus. We also established NRG1’s functional contribution through loss-of-function approaches, and ligand rescue or selective disruption of NRG1–ErbB3 signaling would help clarify the receptor-level mechanism. Finally, in vivo work is needed to test whether alcohol induces NRG1 in mammary epithelium or tumors, and whether blocking NRG1/ErbB3 signaling reduces alcohol-associated tumor promotion.

## Conclusion

In summary, this study identifies NRG1 as a previously unrecognized mediator connecting the estrogenic effects of alcohol with ErbB3/RTK signaling in ER+ breast cancer cells. Our findings support a model in which alcohol activates ER-dependent transcription and induces NRG1, while NRG1-dependent ErbB3 signaling feeds back to reinforce ER activity and downstream proliferative and invasive phenotypes. This mechanism expands the conventional view of alcohol as an estrogenic stimulus by showing how ER activation may be translated into a broader growth-factor signaling network. The ER-NRG1-ErbB3 axis therefore provides a mechanistic framework for understanding how alcohol can exploit signaling circuitry characteristic of ER+ breast cancer and identifies NRG1/ErbB3 signaling as a potentially important interface between a modifiable environmental exposure, tumor progression, and therapeutic response.

## Data Availability Statement

All data is contained within the manuscript.

### Author Contributions

ZM: Data curation and analysis, writing-review & editing; AP: Data curation and analysis, writing-review & editing; XY: Conceptualization, funding acquisition, investigation, project administration, writing – drafting, editing & review. All authors have read and agreed to the published version of the manuscript.

## Acknowledgments

The authors extend their appreciation to the funding agency and support from colleagues in this department.

